# Modeling the consequences of heterogeneity in microbial population dynamics

**DOI:** 10.1101/124412

**Authors:** Helena Herrmann, Conor Lawless

## Abstract

**Chapter 1:** The rate at which unicellular micro-organisms progress through the cell cycle is a major component of their evolutionary fitness. Measuring fitness phenotypes in a given environment or genetic background forms the basis of most quantitative assays of drug sensitivity or genetic interaction, including genome-wide assays. Growth rate is typically measured in bulk cell populations, inoculated with anything from hundreds to millions of cells sampled from purified, isogenic colonies. High-throughput microscopy reveals that striking levels of growth rate heterogeneity arise between isogenic cell lineages (Levy *et al*., 2012). Using published *Saccharomyces cerevisiae* data, I examine the implications for interpreting bulk, population scale growth rate observations, given observed levels of growth rate heterogeneity at the lineage level. I demonstrate that selection between cell lineages with a range of growth rates can give rise to an apparent lag phase at the population level, even in the absence of evidence for a lag phase at the lineage level. My simulations further predict that, given observed levels of heterogeneity, final populations should be dominated by one or a few lineages.

**Chapter 2:** In order to validate and further explore the conclusions from Chapter 1, I re-analyzed high-throughput microscopy experiments carried out on Quantitative Fitness Analysis (QFA) *S. cerevisiae* cultures (Addinall *et al*., 2011), an approach referred to as *μ*QFA. To allow for precise observation of purely clonal lineages including very fast-growing lineages and non-dividing cells, I re-designed an existing image analysis tool for *μ*QFA, now available as an open source Python package. Fast-growing outliers in particular influence the extent of the lag phase apparent at the population level, making the precision of growth rate estimation a key ingredient for successfully simulating population observations. *μ* QFA data include population observations which I used to validate the population simulations generated from individual lineage data. I explored various options for modeling lineage growth curves and for carrying out growth rate parameter inference, and included the full workflow in an open source R package.

## CHAPTER 1: POPULATION-LEVEL OBSERVATIONS FAIL TO CAPTURE ISOGENIC GROWTH RATE HETEROGENEITY

### 1 INTRODUCTION

Cell growth rate, a measure of how quickly cells progress through the cell cycle, is an important component of evolutionary fitness in unicellular micro-organisms and is thus subject to great selective forces. In optimal conditions, microbes with reduced growth rate are rapidly out-competed in bulk populations, making growth rate the ultimate phenotypic indicator of cell health. When modeling cell population dynamics the growth rate parameter is typically measured by making bulk observations at the population scale. Based on the assumption that population growth is representative of individual lineage growth rates, analyses done at the population level are of technical convenience. Population observations, however, ignore growth rate heterogeneity at the clonal lineage level (Van Dijk *et al*., 2015; Kiviet *et al*., 2014).

Modern, automated microscopy and microfluidics offer increasing evidence that, even among isogenic populations, there is considerable heterogeneity in growth rates (Pin and Baranyi, 2006; Schmidt *et al*., 2012; Levy *et al*., 2012). The idea of phenotypic heterogeneity arising through non-genetic differences, such as epigenetics (Bird, 2007) and cell age (Ginovart *et al*., 2011), is beginning to receive much-needed attention as it finds applications in modeling the dynamics of microbial infections, food security assessments, and tumorigenesis dynamics, to name a few. By analyzing the effect of growth rate heterogeneity in clonal *Saccharomyces cerevisiae* lineages, I aim to provide a tractable model for precise measurement of unicellular eukaryote growth rate.

High-throughput, single lineage *S*. *cerevisiae* data allow capturing of heterogeneity between growth rates in isogenic lineages. By fitting simple growth models to single lineage time-lapse data, I simulate population growth from clonal growth rate distributions. I quantify the effects of lineage selection to obtain new mechanistic insight into the early phases of microbial population growth (Figure 1 (i, ii)). Typical growth phases, including the lag and exponential phases of growth, are commonly observed at the population level. Given only population observations it is reasonable to assume that an observed lag phase arises at the single lineage level also.

**Fig. 1:**
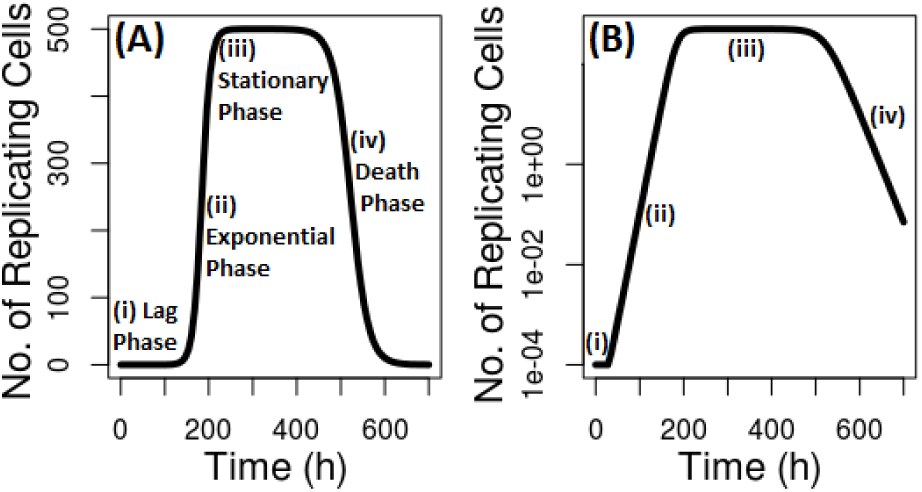
Textbook microbial growth curve: Typical population growth is shown on the original scale (A) and on a logarithmic count (B). Growth phases consist of a (i) lag phase, during which inoculated cells adapt to their new environment, (ii) an exponential growth phase, during which cells divide at a constant growth rate, (iii) a growth arrest phase, during which cell population resources run out (iv) and a death phase, during which viable cell counts are declining.

Observed lag phases are typically associated with the time required for inoculated cells to adapt to their new environment (Rolfe *et al*., 2011). Pirt (1975) however notes that an apparent lag phase may arise as a result of a mixture of growing, non-growing and dying cells in a population, whereby a lag phase can appear in bacterial cultures for up to 20 generations. Defining an apparent lag phase as an observed change from slow to fast growth purely at the population level, I show that surprising levels of heterogeneity in the growth rate distributions as observed in the published data set by Levy *et al*. (2012) can give rise to an apparent lag phase at the population level, without any lag phase at the lineage level. I capture the evolutionary process of cell growth by fitting simple population growth models to thousands of their high-throughput, single lineage *S. cerevisiae* data and by simulating population growth. Simulated population observations confirm that population growth rates are, during the exponential growth phase, often exclusively driven by the fastest growing strains within a population. The effect of fast-growing outliers masking underlying growth rate heterogeneity is especially misleading when, upon induced stress, slow-growing sub-populations provide a selective advantage and subsequent population dynamics are drastically altered, as has been shown to be the case in drug resistance and chemotherapy evasions (Balaban *et al*., 2013; Marusyk *et al*., 2012). Tabassum and Polyak (2015) mark the understanding of the dynamics underlying isogenic, heterogeneous cell lineages as a requirement for developing more effective cures in cancer research.

### 2 DATA SETS & SOFTWARE

I re-analyzed previously published single lineage *S. cervisiae* data from Levy *et al*. (2012) kindly provided by Sasha Levy and Mark Siegal. Lineage data were generated by observing cells inoculated in a layer of Concanavalin A at the bottom of 96-well plates at the single lineage level and using automated microscopy to generate 10 h long time courses with images taken every hour. Captured colony areas were used as a surrogate for cell number. *S. cerevisiae* strains *yme1*Δ, *pet9*Δ, *yfr054c*Δ, *yhr095w*Δ, *snf6*Δ, *rad50*Δ, *not5*Δ, and *htz1*Δ were analyzed.

Some clonal lineages in the provided data contained time courses with missing values. Time courses with few observations can arise as a result of image analysis failures (Levy *et al*., 2012) or fast-growing lineages (microcolonies are tracked only until they touch a neighboring colony). In order to eliminate experimental errors yet not to bias against fast-growing lineages a time course length of three consecutive observations starting from the first observed time point (*t* = 0) was set as a minimum requirement. Growth curves which did not fulfill the latter requirement were discarded from further analyses.

Data analyses and subsequent simulations and visualizations were carried out using R (version 3.3.2; R Core Team, 2016). All computational analyses have been packaged in the detstocgrowth R package available at https://github.com/lwlss/discstoch/tree/master/detstocgrowth.

### 3 METHODS

#### 3.1 Capturing growth dynamics

All growth curves were checked for the existence of a lag phase by assessing linear regression fits on the logarithmic scale. A change in slope on the logarithmic scale marks a change in colony growth rate. Logarithmic area estimates of each growth curve were fed into the bcp R package (version 4.0.0; Erdman *et al*., 2007) which assesses the probability of segmentation in a straight line using a Bayesian approach. A break point probability (*b_p_*) and its location (*b_l_*) are returned.

Growth curves with *b_p_*< 0.5 were modeled using an exponential growth model on the logarithmic scale in the form of a log-linear regression:

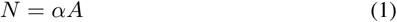

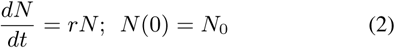

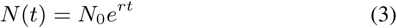

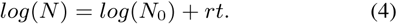

Area *A* is proportional to the number of cells *N* by a conversion factor *α*. Exponential growth of an initial population size *N*_0_ over time *t* is captured by growth rate *r*.

Growth curves with *b_p_* ≥ 0.5 were modeled using a piecewise log-linear regression with the first segment corresponding to the lag phase and the second segment corresponding to the exponential phase. I used the segmented R package (version 1.4; Muggeon, 2003), which fits two continuous straight lines with different slopes to the data. I assumed a single break point with the initial guess being the break point location returned by the bcp output. Using

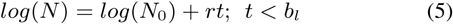

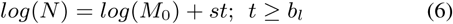

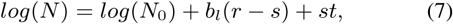
growth rate estimates *r* for the first segment and *s* for the second segment are obtained. *M*_0_ corresponds to the population size at *t* = *b_l_*.

#### 3.2 Parameter inference for single lineages

##### 3.2.1 Inference using a deterministic model

Individual lineage growth curves show no significant evidence for segmentation (*b_p_* < 0.5). Single cell lineages show exponential growth for the duration of the observations and are thus modeled using Equation 4 (mean *R*^2^= 0.94). This differs from most population observations where an observed change in growth rate from the lag to exponential phase as shown in Figure 1 would result in a piece-wise linear growth (Buchanan *et al*., 1997; Baranyi, 2002). A log-normal measurement error model was assumed. The estimated growth rate parameter thus corresponds to the slope of the straight line fitted to the cell density estimates measured in area on the logarithmic scale. Growth rate estimates less than zero were set equal to zero since negative microbial growth holds no biological meaning in this context. I then generated growth rate distributions as implemented in `detstocgrowth` summarizing the growth rate estimates obtained for each strain.

##### 3.2.2 Inference using a stochastic model

In order to assess how inter-lineage stochasticities affect lineage growth, growth rates for the single lineage data were also inferred using a stochastic, birth-only model. I made use of a newly-developed (unpublished) implementation in `Julia` (version 0.4.6; Bezanson *et al*., 2012), kindly provided by Jeremy Revell. Lineage growth is modeled using a Gillespie stochastic simulation algorithm (Gillespie, 1977). Cell division is captured by a single reaction *C* → *2*C, with logistic growth (Verhulst, 1845) propensities, *h*, such that *h_i_* = *C_i_*(1 − 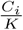) for each cell *C*. The model was run for 1000 samples with *K* = 1000000 and initial growth rate estimate *r* =0.5. Parameter updates follow a cross-entropy method which minimizes the distance between the probability distribution of the data and that of the model (Rubenstein and Kroese, 2004). Strain growth rate distributions were plotted using the obtained parameters as before. Stochastic simulations were compared to deterministic simulations for individual growth curves using parameters inferred from the two models.

#### 3.3 Population simulations using empirical data

Pirt (1975) characterized apparent population growth rate by modeling a dividing and non-dividing population using the 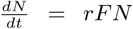 growth equation where *F* refers to the fraction of dividing cells. I modeled the full extent of isogenic heterogeneity by simulating population growth as follows:

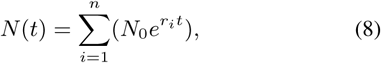
where *r_i_* refers to the growth rates of the *n* lineages in a population. This equation no longer assumes an average population growth rate; instead it captures the growth of each individual lineage.

In order to simulate clonal bulk population growth, I randomly sampled lineage growth rate parameters *r_i_* from growth rate distributions obtained for each strain in the parameter inference step. Individual growth curves were generated according to Equation 3 with a starting population of *N_0_* = 1. Simulated growth curves of single lineages were summed to give the population size at hourly time points over four days using Equation 8 where *N*_0_ = 1 and integer *n* ∈ [100, 10000]. A time course of four days was chosen as it corresponds to the typical duration of a serial dilution experiment capturing population growth such as Quantitative Fitness Analysis (QFA) (Addinall *et al*., 2011). Optimal growing conditions were assumed for the length of the time course. For each strain, this process was iterated 100 times to give mean estimates for population parameters.

As a result of selection, population simulations frequently show a discrete change in growth rate (*b_p_* ≥ 0.5), resulting in a piece-wise linear growth curve on the logarithmic scale (Figure 2). Population growth was thus modeled using Equation 7. Growth rates for both the slow (lag) and fast (exponential) growing phases of the simulated population growth curves were obtained and compared using a paired two-sample Student’s t-test (Welch, 1947) with a confidence level of 0.99. Apparent lag phase duration estimates corresponding to the point of segmentation of the piece-wise regression were obtained for 100 simulated population observations from which the mean apparent lag phase and the 95% confidence interval were calculated.

**Fig. 2:**
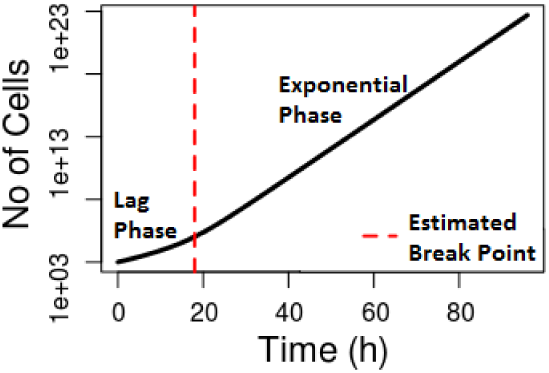
Estimated population growth by fitting a piece-wise log-linear regression: Population growth was simulated according to Equation 8 for *n* = 100 and *N*_0_ = 1, using growth rates sampled from the *yfr054c*Δ distribution. The estimated change point of the fitted piece-wise log linear regression is shown as a red, dashed line.

The *yfr054c*Ɗ strain, which contains the fastest growing lineage in the data set, and the *snf6*Δ strain, which displays a uniquely wide and multimodal growth rate distribution, were chosen as example case studies for assessing the implications of clonal heterogeneity at the population level; calculations for all remaining strains were performed as supporting evidence.

In order to quantify the rate at which lineages expand to dominate bulk populations, percentage contributions of each lineage to a population at a given time as well as the number of strains contributing to more than 5% of the population were averaged for 1000 simulations with population start sizes *n* = 50, 100, 500, 1000, 5000, 10000. Sets of individual example lineages which make up such a population were visualized using the `fishplot` package (version 1.1; Miller, 2016) providing two single-case examples for *n* = 10000 for the *yfr054c*Δ strain.

#### 3.4 Population simulations using synthetic data

In order to further assess the implications of clonal growth rate heterogeneity (captured in the form of distribution width, tail length and multimodality) at the population level, six synthetic lineage growth rate distributions were generated in R, using the LambertW (version 0.6.4; Georg, 2016) package (Figure 3):

1. 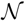 (0:3; 0:015)

2. 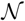 (0:3; 0:015)

3. *t*(0:3; 0:035; 100) with heavy-tail parameter *δ* = 0:08

4. 0:5 * 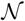 (0:2; 0:03) + 0:5 * 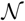 (0:4; 0:03)

5. 0:8 * 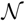 (0:2; 0:03) + 0:2 *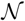 (0:4; 0:03)

6. 6. 0:2 * 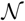 (0:2; 0:03) + 0:8 * 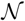 (0:4; 0:03)

**Fig. 3:**
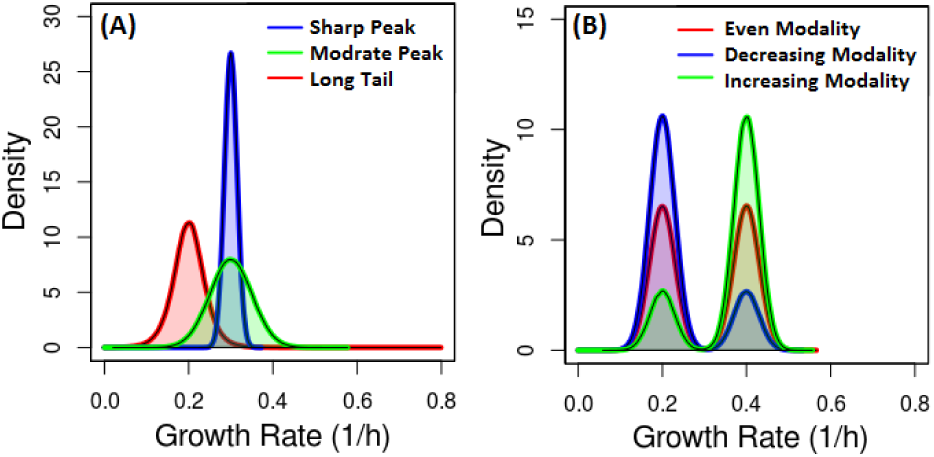
Synthetic unimodal and bimodal lineage growth rate distributions: Three unimodal distributions (A) with different peak heights, widths and tail-lengths and three bimodal distributions (B) with one even and two uneven bimodalities were compared.

Population growth was then simulated from all six synthetic distributions as described in Section 3.3.

### 4 RESULTS

#### 4.1 Population simulations show a piece-wise linear fit on the logarithmic scale in the absence of a lag phase in single lineages

Growth rate parameter inference shows that single lineage growth rates within all strains show remarkable heterogeneity (Figure 4), signified not only by the width of the distributions but also their modality. While the *snf6*Δ strain shows a curioustrimodality (A), a significant fraction of non-dividing and outlying, fast-growing lineages can be found within all strains, resulting in bimodal long-tailed distributions (B).

**Fig. 4:**
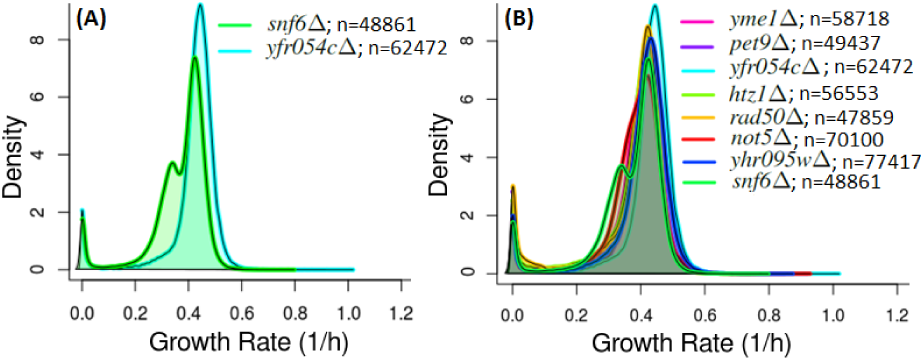
Quantifiable growth rate heterogeneity between isogenic *S*. *cervisiae* lineages: Growth rate distributions for *snf6*Δ and *yfr054c*Δ (A) are overlayed on the growth rate distributions of all published strains (B) in the data set. Distributions were obtained by fitting a log-linear regression to all n single-lineage time courses of each strain.

Figure 5 displays the effects of lineage heterogeneities at the population level for *yfr054c*Δ and *snf6*Δ for inocula *n* = 100, 1000, 10000. Remarkably, although individual lineages are simulated according to a linear fit on the logarithmic scale, population growth curves obtained from Equation 8 are characterized by piece-wise linear growth on the logarithmic scale, a feature which is often assumed to be the result of cell size expansion in preparation for growth (Rolfe *et al*., 2011).

**Fig. 5:**
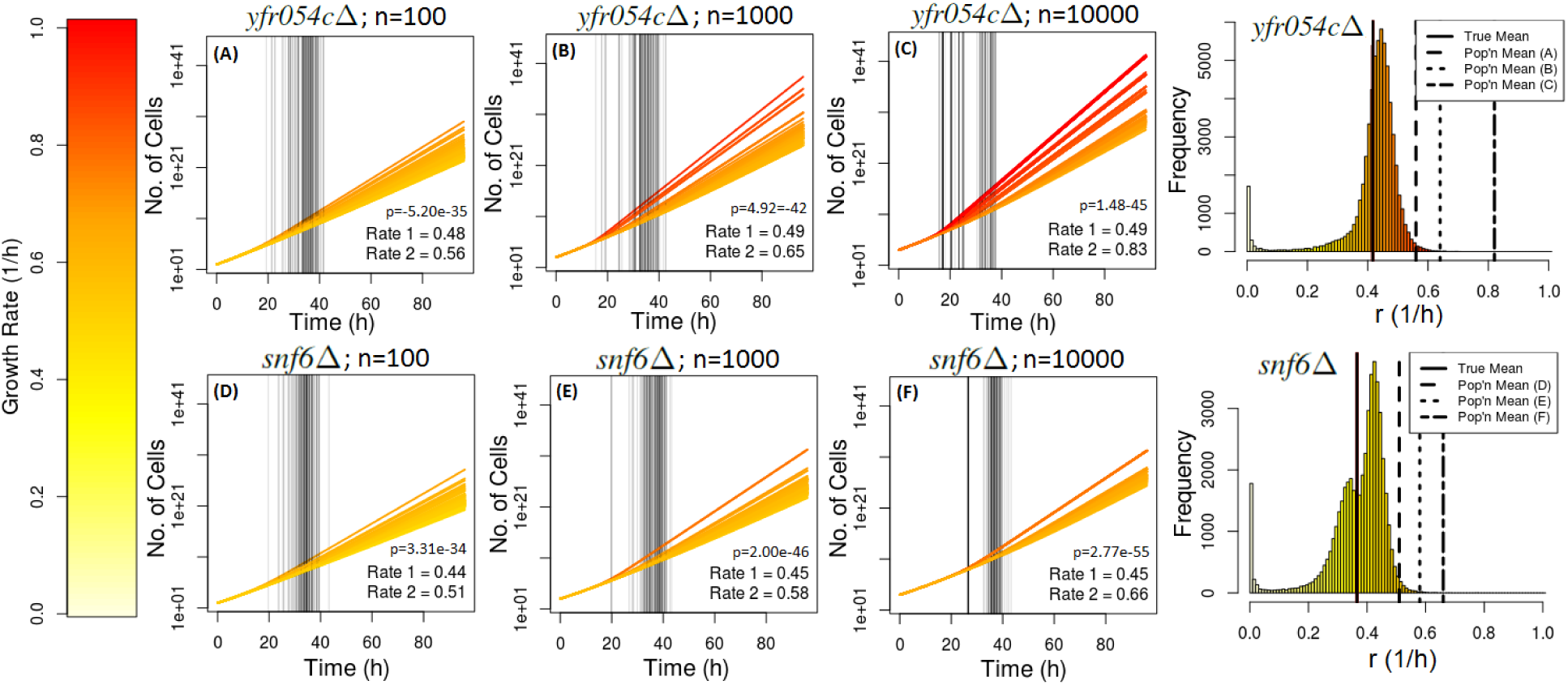
Population simulations from single lineage data show an apparent lag phase: Population simulations over a time course of 1-96 h using Equation 8 for *yfr054c*Δ and *snf6*Δ where *n* = 100, 1000, 10000 growth rates *r_i_* were sampled from the distributions as respectively indicated on the right are displayed. Mean simulation growth rates of the exponential phases of (A-F) are drawn on the distributions. Population simulations are color-coded according to the fastest strain sampled; the color scale displayed on the left applies to all figures. Growth rate parameters were inferred using the same method as in Figure 4. Break point locations for *bp* ≥ 0:5 are marked by vertical black lines. Mean growth rate values for the lag (Rate 1) and exponential phase (Rate 2) are shown along with the significance values obtained from respective Student’s t-tests.

Simulated population growth is significantly greater than mean lineage growth rate and is driven by the fastest sampled strain (Figure 5). The longer tail in the *yfr054c*Δ distribution results in a faster population growth than that observed in the *snf6*Δ simulations. Estimated break points for *b_p_* ≥ 0.5 are marked as vertical lines dividing population growth into a slow-growing (lag) and a fast-growing (exponential) phase. Although the difference between mean lag and mean exponential growth is greatest for *yfr054c*Δ (C), paired two-sample Student’s t-tests reveal the differences between lag and exponential growth to be most significant for *snf6Δ* (F). Break point location is most variable for the *yfr054cΔ* strain which has the widest-ranging growth rate distribution (Figure 4). Population simulations for all strains display an average *b_p_* of 0.92 (minimum *b_p_* = 0.43) and, on average, show a significant change in growth rate for integer *n* = [100, 10000] inocula 99% of the time. For all growth curves with *b_p_* ≥ 0.5, mean growth rates of the slow-growing phase are significantly slower than growth rates of the fast-growing phase (paired two-sample Student’s t-test; *p* < 0.01).

#### 4.2 Observed population growth rates increase with inoculation size and significantly surpass average observed growth rates of single lineages

As previously shown, observed population growth rates during the fast-growing phase correspond to the right-hand tail of the single lineage distributions (Figure 5). Figure 6 predicts a sharp increase in observed growth rate with inoculation size which then begins to level off. Growth rate averages for both segments are significantly greater than the mean of the growth rate distributions. The second segment of the piece-wise linear regression levels off after a greater inoculation size than the first segment, as it is more greatly affected by the variability of sampling fast-growing lineages. The peculiar increase in variance with inocula size is the result of discontinuities in the tail of the growth rate distributions. As inoculum size increases, the probability of sampling from the tail of the distribution increases. Growth rates sampled from the right-hand tail have the largest effect on variance. Figure 6 shows that the number of fast simulations increases with inoculum size. Variance increases are greater for *yfr054c*Δ whose growth rate distribution contains more fast-growing lineages. Evidently, by definition of the Central Limit Theorem (Rice, 1995), variance will approach zero as inoculum size approaches infinity; this however does not apply to the inocula sizes considered here, where starting population sizes are smaller than the number of observations in the distributions from which growth rates are sampled.

**Fig. 6:**
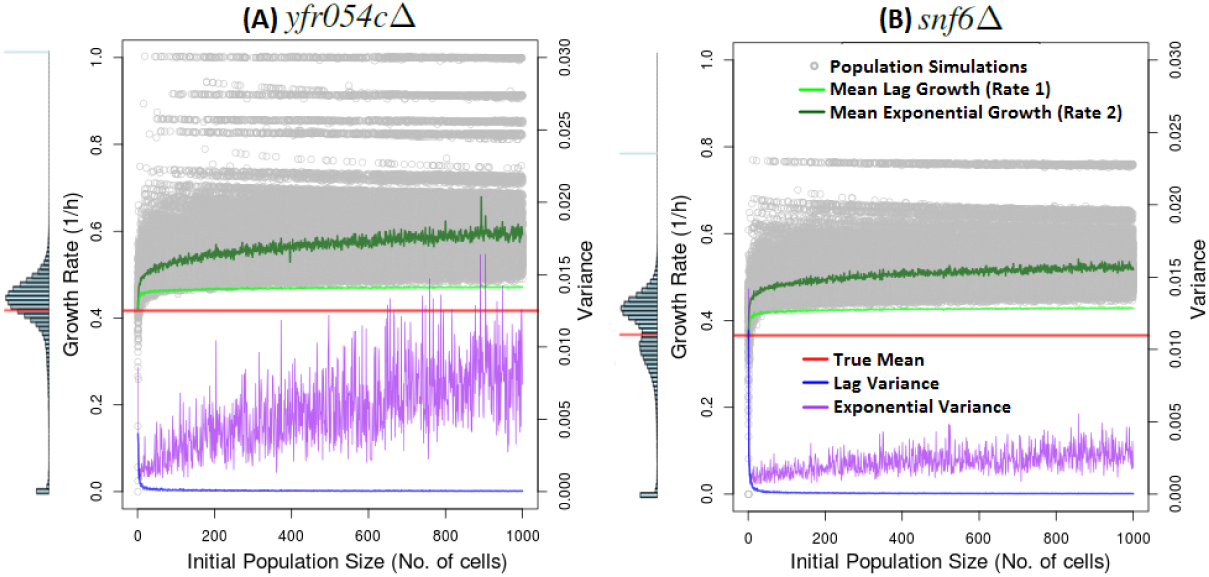
Population growth rate estimates and reproducibility with increasing inoculation size: Growth rates of *yfr054c*Δ (A) and *snf6*Δ (B) populations inoculated with 1-1000 cells (grey) were computed. The mean and variance of 100 iterations of the two segments of a piece-wise linear regression were calculated and averages for each population start size are displayed (light green and blue for the lag phase; dark green and purple for the exponential phase). The mean lineage growth rates of the two distributions shown as vertical histograms on the left of each figure are marked in red. Histograms are on the same growth rate scale as the main figure.

#### 4.3 The duration of the apparent lag phase of the population simulations decreases with inoculation size

Mean apparent lag phase duration predicts a negative relationship between inoculation size and observed lag phase (Figure 7). A longer tail in the growth rate distribution of *yfr054c*Δ is suspected to increase the effect of selection for fast growers, resulting in an earlier onset of the exponential phase as compared to *snf6*Δ. For *n* = 100, samples from the right-hand tail of the distributions are improbable; thus lag phase estimates of *yfr054c*Δ and *snf6*Δ match closely, since the central peaks of the growth rate distributions overlay (Figure 4).

**Fig. 7:**
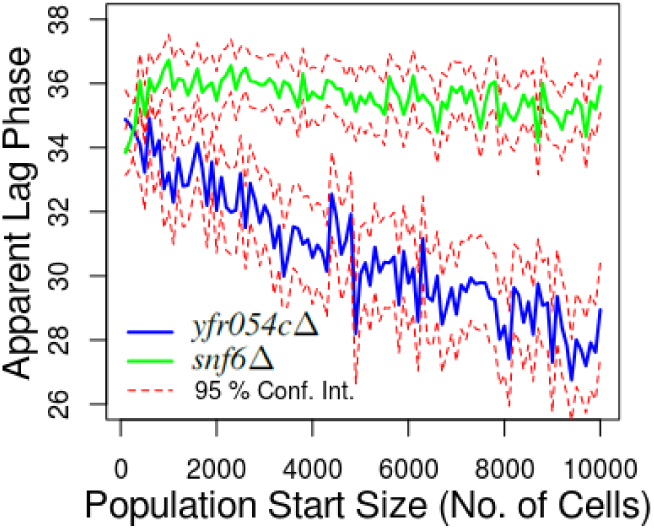
Break point estimates over time in simulations of *yfr054c*Δ and *snf6*Δ: Apparent lag phase durations obtained by fitting Equation 7 to the population simulations generated using Equation 8. Mean estimates along with their 95% confidence intervals (Conf. Int.) of 100 iterations are shown for integer population sizes *n* = [100, 10000] in steps of 100.

#### 4.4 The lag phase apparent at the population level is the result of selection between lineages with heterogeneous growth rates

Figures 5 and 6 suggest that a selection process acting on heterogeneous isogenic lineages drives the observed population growth rate to be significantly faster than the average growth rate of single lineages. This is confirmed by looking at the lineage compositions (Figure 8). Population simulations of *yfr054c*Δ show how the fastest sampled strains dominate the population over time and drive exponential population growth rate. The lag phase apparent at the population level is thus the direct result of strong lineage selection.

**Fig. 8:**
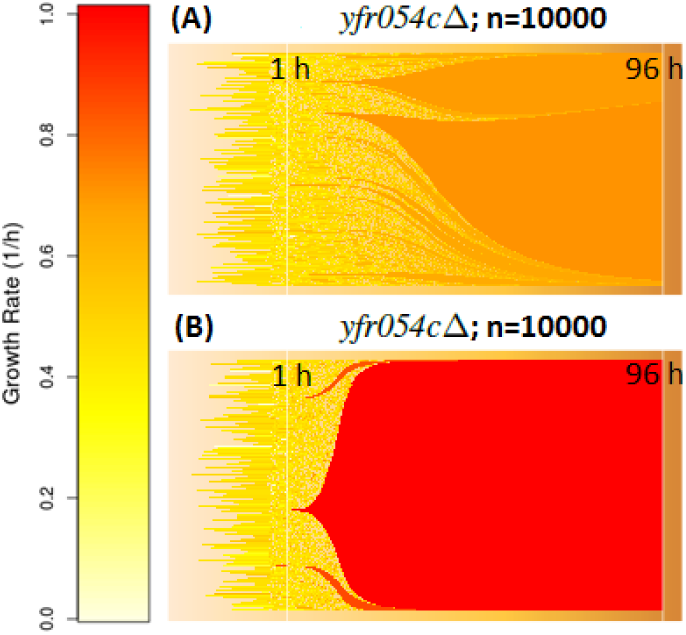
Lineage composition of population simulations highlight an extensive selection process: Individual lineages which make up two population simulations of *yfr054c*Δ shown in Figure 5 (C). Each horizontal line corresponds to a single clonal lineage. Percentile population compositions of clonal lineages from 1-96 h are shown.

The distinction between (A) and (B) further highlights how variability in population growth rate is affected by the sampling of fast-growing lineages. Faster-growing lineages decrease the duration of the apparent lag in population growth as they dominate the population more quickly.

Mean dominance effects of single fast-growing lineages within a population (Figure 9) confirm that, once they make up a significant percentage of the population, individual fast-growing lineages drive a change from slow to fast growth at the population level. On average, less than five lineages make up more than 5% of population each. Figure 9 predicts that in *yfr054c*Δ populations simulations, the single fastest lineage is, on average, slightly more dominant than those of *snf6*Δ. The length of the right-hand tail of the lineage growth rate distributions thus determines the extent of selection and the lag phase apparent at the population level. Increased inoculum size results in greater dominance of fast-growing strains since the probability of sampling fast-growing lineages is higher (Figure 9).

**Fig. 9:**
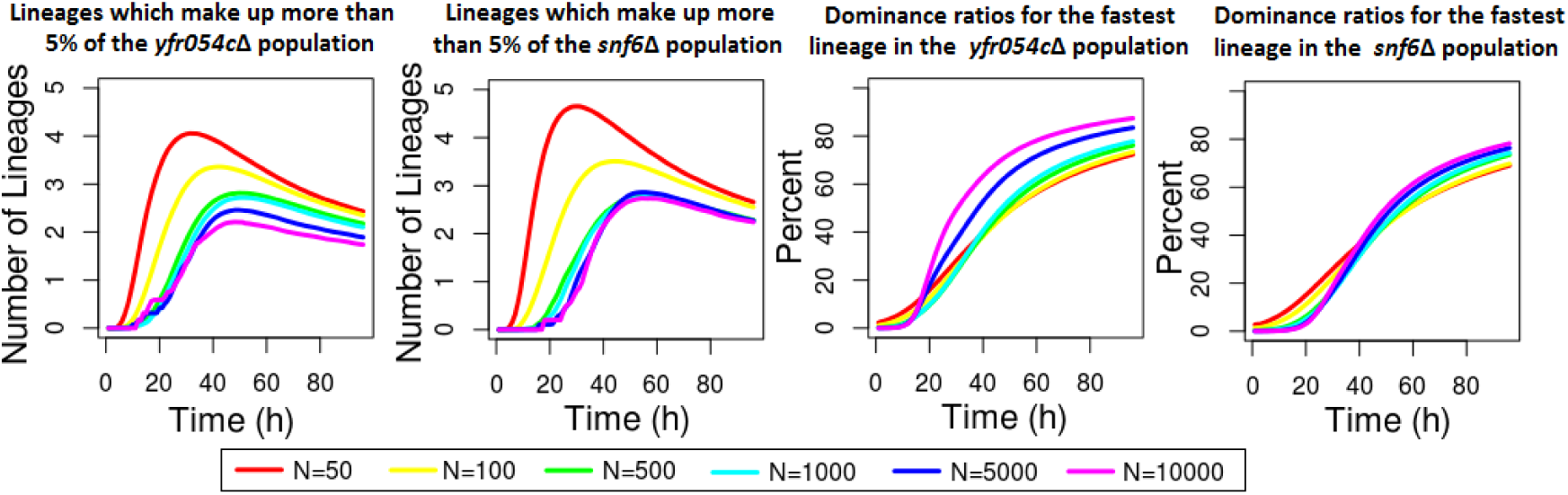
Dynamics of population structure in simulated selection are affected by inoculum density: The mean number of simulated lineages which make up more than 5% of the population over time (left) and the mean percentage with which the simulated lineages of the fastest sampled growth rate make up the population independent of the number of times that particular growth rate was sampled (right) are shown. All trials show the average of 1000 simulations for the *yfr054c*Δ and *snf6*Δ strains where populations are simulated using Equation 8 for *n* = 50, 100, 500, 1000, 5000,10000 from 1-96 h.

#### 4.5 Simulated selection between synthetic lineages confirms that the width and tail-length of the growth rate distribution influence the apparent population behavior

Heterogeneity at the single lineage level influences observations made at the population level. While Figures 4-9 are based on true experimental data, the results shown in Figure 10 make use of synthetic distributions in order to quantify how features of heterogeneity at the single-lineage level, as captured by the shape of the distribution, affect population level observations. A sharp peak in the growth rate distribution results in some evidence for an apparent lag phase (A); the observed change in growth rate is significant (*b_p_* ≥ 0.5) in 96% of the simulations. Widening the distribution increases the extent of the apparent lag phase with all population simulations showing a significant change in growth rate due to an increase in average population growth rate during the exponential phase (B). If the right-hand tail of the growth rate distribution is widened, the variability in population simulations increases;this again emphasizes that population behavior is heavily influenced by the fastest strain selected (C). Thus, greater growth rate heterogeneity at the lineage level in the form of a wide or long-tailed growth rate distribution increases the extent (increased *b_p_*, greater difference in growth rate between phases and decreased duration) of the apparent lag phase. Figures 10 (D-F) show that modality *per se* has little effect on the observed population behavior, implying that slow-growing sub-populations and non-dividing cells are masked by population level observations. However, increasing the number of fast-growing lineages without lengthening the tail of the distribution extends the apparent lag phase (D, F). A lower number of fast-growing lineages increases the variability in observed exponential growth rate (C, E). Even modality leads to an increased variability in lag duration (D).

**Fig. 10:**
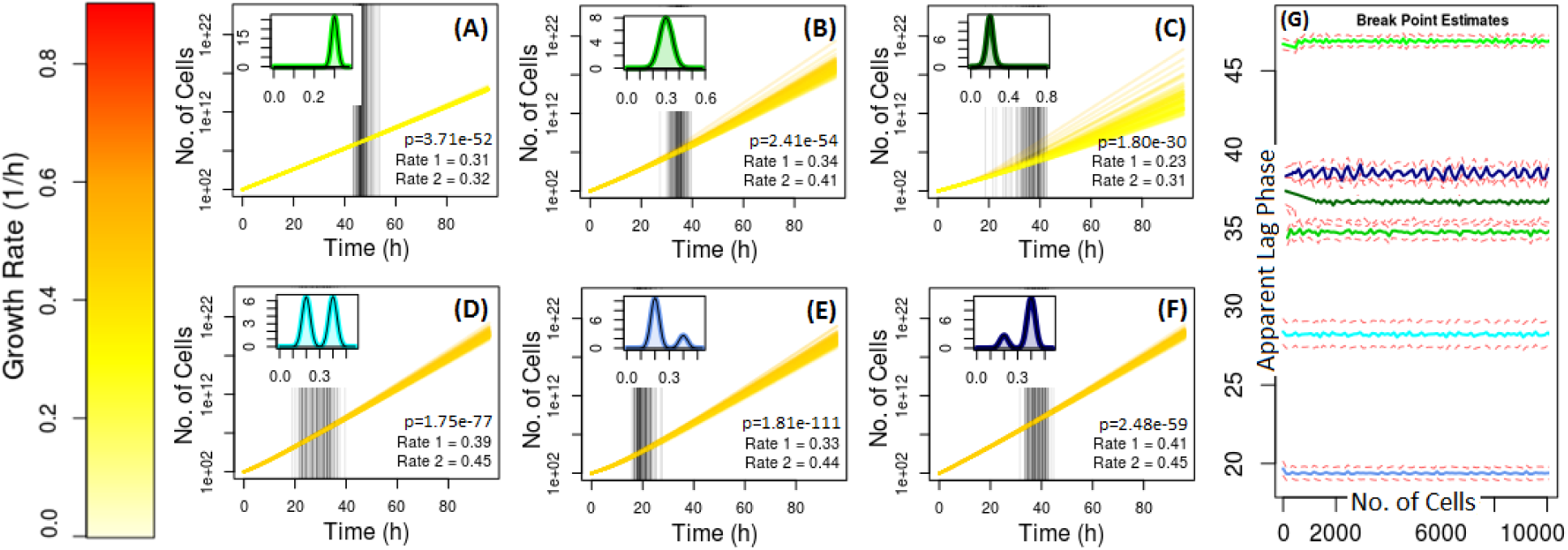
Population simulations obtained from synthetic lineage growth rate distributions: Population simulations as shown in Figure 5 based on generated synthetic distributions (Figure 3) are shown. A miniature version of each growth rate distribution from which the respective population simulations were generated is shown in the top left (A-F). Estimated break points (*b_p_* ≥ 0.5) are again marked by dashed, vertical lines. Predicted lag phase durations for integer population start sizes *n* = [100, 10000] are displayed on the right, color-coded according the distributions shown in (A-F).

The summary of lag duration (G) shows that lag duration decreases as the distance between lag and exponential growth increases as a result of fast-growing outliers. An observed decrease in lag duration as a result of increased number of fast-growing lineages with increased inoculation size shown in Figures 6 and 7 is confirmed for the first 1000 cells when sampling from a long-tailed distribution (C). Continuity in the distribution here causes the decrease in lag to level off faster than observed in Figure 7. Similarly, as the sub-population of fast-growing lineages becomes greater than the sub-population of slow-growing lineages (F), lag duration is increased and vice-versa (E). Increasing the fast-growing sub-population (F) increases the probability of sampling fast-growing lineages and thus increases the observed variability in lag duration (G).

#### 4.6 Accounting for inter-lineage heterogeneities affects individual growth rate estimates

When modeling lineage growth using a stochastic, birth-only model, obtained growth rate parameters can differ from those obtained when fitting a log-linear regression (Figure 11). Strain distributions overlay closely; this suggests that interlineage heterogeneities are marginal and have limited effect on observed lineage growth rate within a strain. However, when looking at individual lineages, growth rate parameters obtained from the two models can differ by up to 0.27 h^−1^ (Figure 11 (A.ii)). Fast-growing outliers such as growth curve 2057 in the *yfr054c*Δ data and growth curve 1901 in the *snf6*Δ data (Figure 11 (A.iii, B.iii)) have previously been shown to significantly influence observed population behavior (Figures 5, 6, 8, 9, 10). Precision in estimating fast-growing outliers is thus of great significance. Figure 11 (B) further suggests that the tri-modality observed in, *snf6Δ* when modeling lineage growth using Equation 4 is an artifact of inter-lineage stochasticities. Also, the multi-modality resulting from a significant number of non-dividing growth curves in the distribution is less distinct for parameters inferred using the stochastic model. This however is due to the fact that the current stochastic implementation assumes growth to be positive and is why population growth was simulated using the parameters obtained from the deterministic model (Equation (4)) as shown previously. The growth rate parameters for nondividing cells are merely approaching zero (Figure 11 (A.i, B.i)) in the stochastic implementation.

**Fig. 11:**
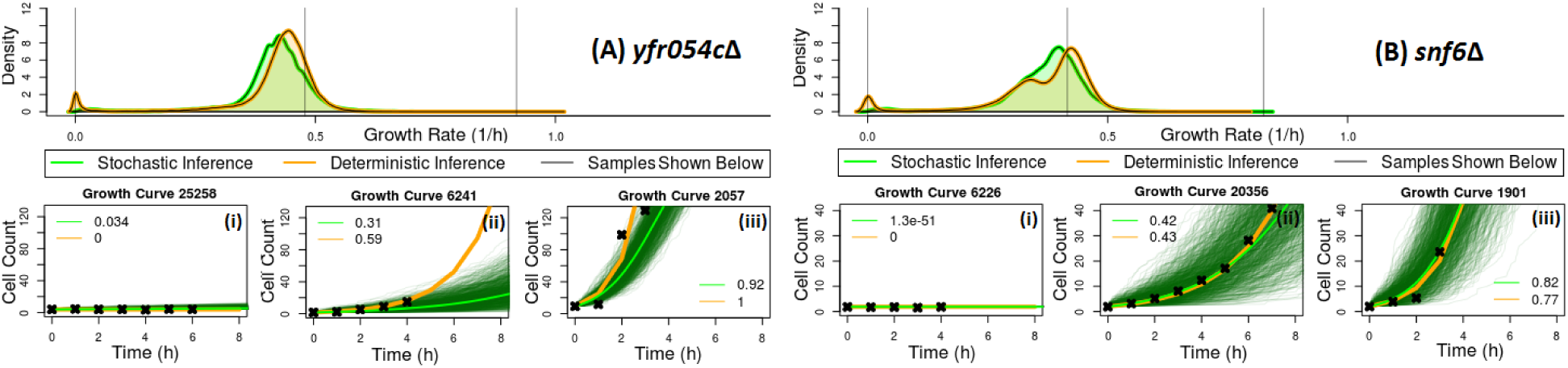
Stochastic and deterministic models of single lineage growth predict slightly different growth rate parameters: Growth rate distributions obtained from deterministic (orange) and stochastic (green) parameter inference for *yfr054c*Δ (A) and *snf6*Δ (B) are compared. Growth curves observations associated with the minimum (i), the maximum (iii) and an estimate growth rate parameter sampled from the central peak of the distribution (ii) obtained using the stochastic model are shown below. 10000 stochastic simulations (dark green) and the deterministic solution (orange) to (Equation 8) with *r* set to the respective inferred growth rate parameter are overlayed. The mean of all shown stochastic simulations is displayed in green.

### 5 DISCUSSION

Population measurements are the most common tool for analyzing strain growth rate as a measure of fitness. I reinterpreted previously observed growth rate heterogeneity among isogenic *S. cerevisiae* populations to examine consequences at the population level. Apparent lag, which arises purely as a result of clonal heterogeneity, is here distinguished from true lag, defined as the time required for inoculated cells to adapt to their new environment (Rolfe *et al*., 2011). While the vast range of observed microbial lag phase patterns has been attributed to many intracellular mechanisms (Rolfe *et al*., 2011) and extracellular stresses (Baranyi, 1998; Fridman *et al*., 2014), this is the first time that, using real data, single lineage heterogeneities are explored as an underlying cause for an observed change in growth rate behavior at the population level.

Population simulations of selection obtained from synthetic distributions confirm a positive relation between significance of apparent lag and lineage heterogeneity. Apparent lag duration and the subsequent change in growth rate are particularly sensitive to fast-growing outliers. This should be kept in mind when removing apparent outliers from analyzed data sets; unless these are confirmed experimental errors, trimming outliers will significantly alter population inferences. Simulations based on both real and synthetic data have shown that informative growth rate phenotype characteristics such as the number of non-dividing lineages available at the single lineage level cannot be captured by population observations. Indeed, clonal heterogeneity has in the recent literature led to new insights on population evolution, particularly in cases where slow-growing lineages provide a selective advantage (Van Dijk *et al*., 2015; Ding *et al*., 2012; Levy *et al*., 2012). Single-lineage growth rate heterogeneity should be considered an integral part of a strain’s phenotype when using growth rate as a measure of fitness.

Observed population behavior not only masks heterogeneity but is also not representative of typical single cell behavior. Having demonstrated how an apparent lag phase can arise purely at the population level, my results provide a key new mechanistic insight applicable to predictive microbiology growth models. Robinson *et al*. (2001), for example, presume that average single lineage lag must be greater than observed population lag, since fast-growing lineage begin to divide earlier. I however show that the apparent lag phase observed at the population level does not translate down to the single lineage level but instead corresponds to the time required for the fastest-growing lineages to dominate the population, at which point the population behavior switches from seemingly slow to fast growth. During exponential growth, population level observations are analyzing only the behavior of a small proportion of lineages favored by selection, making the observed population growth rate significantly faster than the mean growth rate of clonal lineages, for example. I suggest modality, modal width and tail-length of the growth rate distributions as more informative measures to summarize strain growth at the single-lineage level.

The selective dominance of a few fast-growing lineages is most prominent in large population sizes and during the exponential phase, agreeing with common biological practices to sample from the exponential phase and to use large starting populations to increase reproducibility (Greenwood, 2012; Jasmin and Zeyl, 2012). However, this only applies for inocula sizes large enough to capture the full range of lineage heterogeneity within a strain. Furthermore, increased reproducibility comes at the cost of accurately describing the whole population of single lineages. Both Van Dijk *et al*. (2015) and Levy *et al*.(2012), for example, consider single-lineage dynamics a necessary step in their analyses for assessing how the competitive dynamics in a population change as a result of induced stress. My findings agree with those of Fridman *et al*. (2014) who show that for strains grown under optimal growth conditions, no lag phase can be observed at the single lineage level and that, at the single lineage level, a lag phase arises as an adaptive response only to antibiotic exposure. In accordance with my simulations, Ginovart *et al*. (2011) observe a decrease in lag phase duration with increased inoculation size up to 1000 *S. cerevisae* cells. An increase in lag phase duration as a result of stress is also a common observation in the literature (Augustin *et al*., 2000; Robinson *et al*., 2001). Under induced stress, slow-growing lineages can outperform fast-growing lineages within microbial populations (Levy *et al*., 2012; Batchelor *et al*., 1997); thus, apparent lag phase observations which arise as a result of selection purely for fast-growing lineages may not apply.

The effect of how stress, including a nutrient and space limiting induced carrying capacity, applied at the single lineage level translates to the population level remains yet to be addressed. Further model limitations include the assumption that cell lineages grow independently. The general shape of the growth rate distributions obtained using a deterministic, exponential model on the log scale was validated using a stochastic model. When comparing the distribution shapes, evidence for inherent stochasticity exists to some extent. For example, bimodalities observed in the center of lineage growth rate distributions, as is the case for *snf6Δ*, are suspected the be the result of inter-lineage heterogeneities. Even though inference of stochastic and deterministic models yields growth rate distributions that match very closely, slight imprecision in growth rate estimates for fast-growing lineages can alter simulated population growth significantly; thus, stochasticity should at least be taken into consideration when estimating single lineage growth rates of future data sets.

Lastly, further investigations are required to determine the origin of heterogeneity. Jasmin and Zeyl (2012) confirm that cells isolated from a population of *S. cerevisiae* over time show a general increase in growth rate as a result of selection for fast growers. However, given that fast-growing lineages dominate population growth and individual cells are always sampled from a population, when does heterogeneity arise? Experiments tracking the heritability of growth rate are yet to be conducted at the single lineage or the single cell level.

## CHAPTER 2: *μ*QFA: DIRECT COMPARISON OF POPULATION SIMULATIONS BASED ON SINGLE LINEAGE DATA WITH TRUE POPULATION OBSERVATIONS

### 1 INTRODUCTION

Chapter 1 effectively outlines the implications of heterogeneity between individual lineage growth rates by simulating bulk cell populations. While the previous analyses were based on published isogenic lineage area estimates over time, I did not have access to the raw data, including colony images. In order to test my population simulations against true population observations, I re-analyzed high-throughput microscopy experiments on Quantitative Fitness Analysis (QFA) (Addinall et al., 2011) cultures in order to observe single lineage growth, an approach I here refer to as *μ*QFA. I re-developed an existing image analysis tool, now available as a Python package, for obtaining cell density estimates. By observing multiple lineages derived from single *S. cerevisiae* cells grown on solid agar plates using automated microscopy, *μ* QFA enables high-content insight into the population observations of standard QFA. High-content insight includes the number of non-dividing cells and the microscopic observations for fast-growing outliers, which as shown in Chapter 1 significantly influence population observations. A single fast-growing lineage has been shown to dominate an entire population (Figure 8 (B)).

The aim here is to more easily distinguish inherent noise from experimental errors in order to obtain more exact growth rate distributions and to validate previous approaches. Population simulations from *μ* QFA lineages are compared to the true population growth observed for each of the observations made under the microscope. This direct comparison between simulated population observations based on single lineage data and true population observations made from the same data is used to test whether the newly obtained high-throughput data of single cell yeast lineages is matched with the appropriate level of mathematical complexity and biological interpretation to model yeast cell growth. I again test the fit of an exponential growth model against the data, and compare growth rate parameters obtained using a frequentist approach with a Bayesian approach. Bayesian inference allows for incorporating biological constraints in the form of prior distributions and returns a probability estimate rather than a single point estimate (Christensen *et al*., 2011). Additionally, I assess the practicality of doing Bayesian parameter inference using a discrete stochastic growth model on a selection of growth curves.

### 2 DATA SETS & SOFTWARE

Laboratory work was carried out in the Institute for Cell and Molecular Biosciences at Newcastle University with the support of Prof. Lydall’s research group. I implemented an image analysis tool for the obtained microscopic observations in Python (version 2.7.6; Van Rossum, 1995), now available as an open source package called `muqfatc` (https://github.com/lwlss/discstoch/tree/master/muqfatc). For the subsequent analyses I again used the `detstocgrowth` package (https://github.com/lwlss/discstoch/tree/master/detstocgrowth) written for the analyses done in Chapter 1, implementing additional features where required.

### 3 METHODS

#### 3.1 *μ*QFA

*S. cerevisiae his3*Δ and *htz1*Δ strains, grown on solid agar plates as part of a library of the Stanford yeast knockout collection, were picked and streaked for single colonies onto solid agar in round Petri dishes. *his3*Δ acts as a wild-type surrogate (it has the HIS3 gene inserted elsewhere in its genome), and *htz1*Δ was chosen for comparison since its growth rates have previously been observed to be heterogeneous (Levy *et al*., 2012). Individual colonies assumed to be derived from a single cell were picked using sterile toothpicks and inoculated into 200 μl volumes in a rectangular 96-well microtitre Nunc plate. Cells were grown to saturation over 48 h and were transferred onto a solid agar surface inside a rectangular plate following the manual QFA procedure described by Banks *et al*. (2012), using a sterile pin tool by V&P scientific. The plate was sealed off with electrical insulation tape and placed under a Nikon Eclipse 50i microscope mounted with a fully automated Prior Optiscan II stage and a Jenoptik ProgRes MF scientific camera. Pin time course images were captured by automated microscopy using the *μ* Manager software (version 1.4.21; Edelstein *et al*., 2010) so that each pin was captured every 20 min over 48 h.

#### 3.2 Time Course Generation

Raw microscopic images were analyzed in Python using the OpenCV package (version 3.0.0; Bradski and Kaehler, 2008). Color images were converted to gray-scale. I then used a Canny edge detection algorithm (Canny, 1986) to capture yeast colony contours. Areas were estimated by counting the number of pixels corresponding to yeast (white) on agar (black) at each time interval using graphic erosion and dilation to smooth edges (Figure 12). Area estimates are based on the assumption that cell growth is two-dimensional; a valid assumption to make at early time points.

**Fig. 12:**
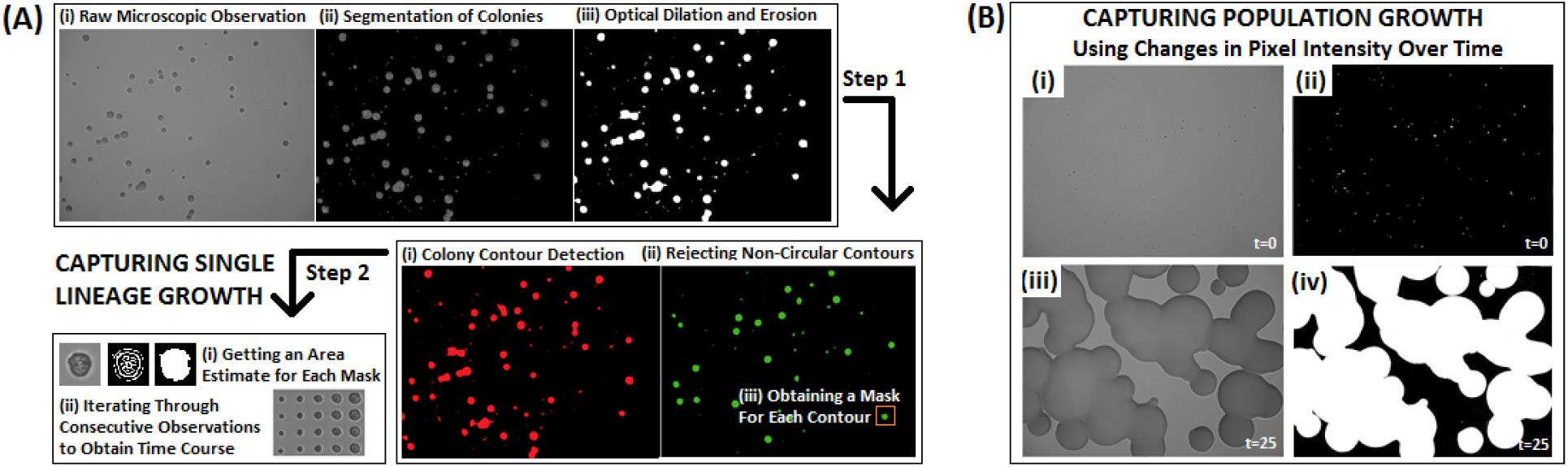
Image analysis work flows for capturing colony growth over time: Upon image segmentation area estimates acting as surrogates for colony density are captured by counting the number of white (yeast) pixels on a black background (agar) over time. Lineage growth rates are estimated using an optical dilation and erosion process and subsequent circular contour detection (A). Colony masks obtained from the final time point are tracked backwards through time to obtain a full time course. Population growth rates are estimated by capturing the total number of pixels corresponding to yeast cells in each pin over time (B). A chosen gray-scale value is used to distinguish yeast from agar.

Colony contours of non-merged colonies were tracked individually for 6 h (20 observations) and checked for circularity comparing the minimum enclosing area to the total area of a colony at each time point to give dynamic, isogenic lineage estimates. After checking growth rate distributions for a range of time course lengths, I chose to set a time course duration of 6 h since this maximizes the number of obtained lineage observations without biasing against fast-growing lineages.

Population estimates were made by segmenting the image according to pixel intensity. Dark yeast colonies were observed on a light agar surface; yeast area estimates correspond to gray-scale values 110 or darker. Population area is estimated up to 28 h for each time course, after which the lighting of the microscopic observations is too dark to distinguish between cells and agar.

#### 3.3 Parameter Inference

Growth rate parameter inference approaches for single lineage time courses are outlined in Figure 13. Full example parameter inference work flows based on the original data are outlined on https://github.com/lwlss/discstoch.

**Fig. 13:**
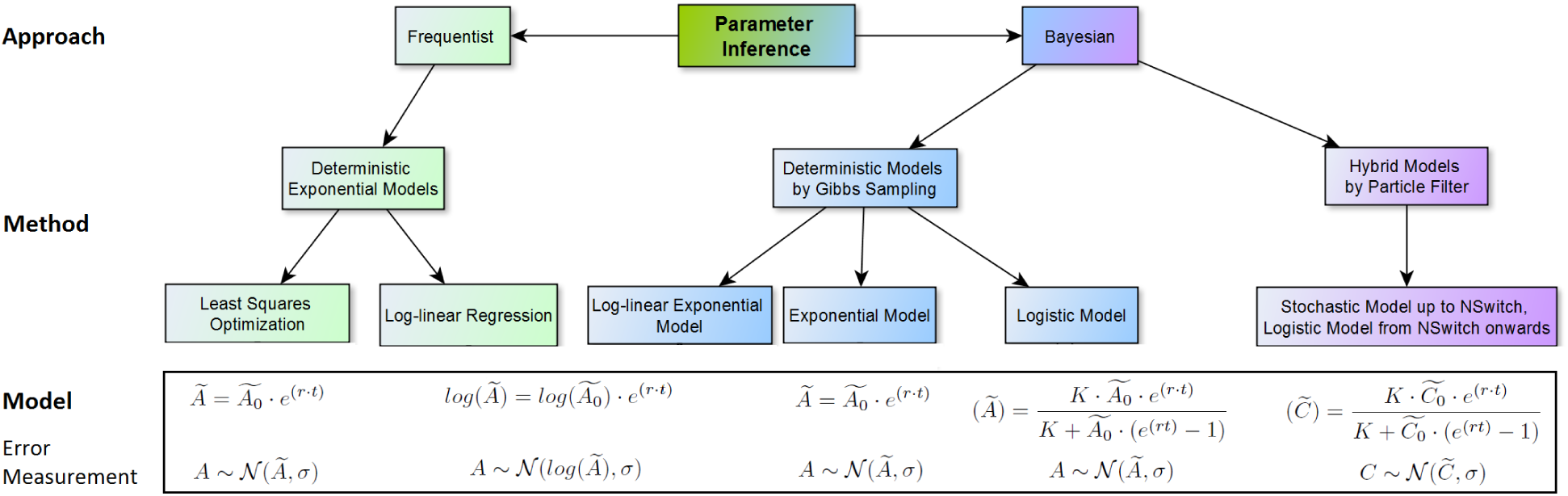
Modeling and inference options assessed for their effectiveness in capturing *μ* QFA single lineage growth rates: Frequentist parameter inference options are displayed in green; Bayesian parameter inference approaches are marked in blue (deterministic modeling) and purple (stochastic & deterministic modeling combined).

I carried out Bayesian inference for deterministic models by Gibbs Sampling using the `rjags` package (version 4.1.0; Plummer, 2016) along with an exponential growth model implementation. Prior distributions for growth rate *r*, initial area *x*_o_ and precision *τ* were specified as follows:

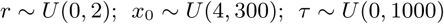

These priors ensure that *r* accommodates a range of growth rates wider than obtained from the deterministic model and that *x*_o_ lies in between the range of areas observed for the first time point. Given 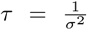,the prior distribution for *τ* accommodates for the minimum standard deviation, *σ* =032, observed among lineage residuals when applying a deterministic model. Resulting growth rate parameter estimates are based on 1000000 iterations with a thinning of 10000. For the implementation of the stochastic model, the `smfsb` R package (version 1.1; Wilkinson, 2013) was adapted to simulate lineage growth according to a stochastic Gillespie, birth-only model (Gillespie, 1977; Bailey, 1964) up to 1000 cells, at which point the variation between stochastic simulations becomes negligible. Subsequent lineage growth was then simulated according to the deterministic logistic growth model (Verhulst, 1845) with a constant carrying capacity *K* = 15000 and initial growth rate estimate *r* = 0.3. (Figure 13). This stochastic/deterministic hybrid implementation was validated using a deterministic (logistic) implementation in rjags with prior distributions as outlined above where identical growth rate estimates were obtained using *K*ϵ [15000, 100000, 1000000]. For the stochastic model I assumed a constant normal error measurement model with *σ* = 20. A single chain was run for a minimum of 1000000 iterations with a thinning of 1000. A tuning parameter of 0.02 was used to update the Markov chain (MC) as implemented in smfsb. In order to convert area estimates into estimates of cell number I assumed

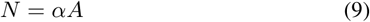

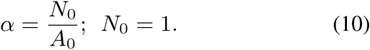

Thus, the maximum likelihood of the distribution of pixel area observed at *t* = 0 can be used as a reciprocal estimate for *α*, such that 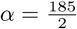.

#### 3.4 Population Simulations

The same procedure as outlined in Chapter 1 was applied, with the time course length set to 48 h rather than 96 h to match the length of this specific *μ*QFA. Population simulations obtained from the growth rates of lineages in the same pins were compared to the total area estimates obtained for the entire pin, allowing me to verify my predictions. Growth curves of the pin population level observations were again modeled using a piece-wise linear regression on the logarithmic scale. Here, however, two break point estimates were used, as the data shows both an apparent lag phase and an apparent carrying capacity. Observations and simulations from row 5, column 5 (counting from the top left-hand corner of the agar plate), denoted as *R*05*C*05, are presented as an example in the results; its microscopic images are well-focused, contain lots of individual growth curves, and are unaffected by possible edge effects which may occur at the side of the plate.

### 4 RESULTS

#### 4.1 Growth rate estimates for clonal *μ* QFA lineages are most accurately inferred using an exponential growth model on the original scale as compared to a log-linear regression analysis

Small colony images are most difficult to capture precisely at the image analysis stage, since blurry microscopic observations display the greatest noise at early time points (Figure 14).

**Fig. 14:**
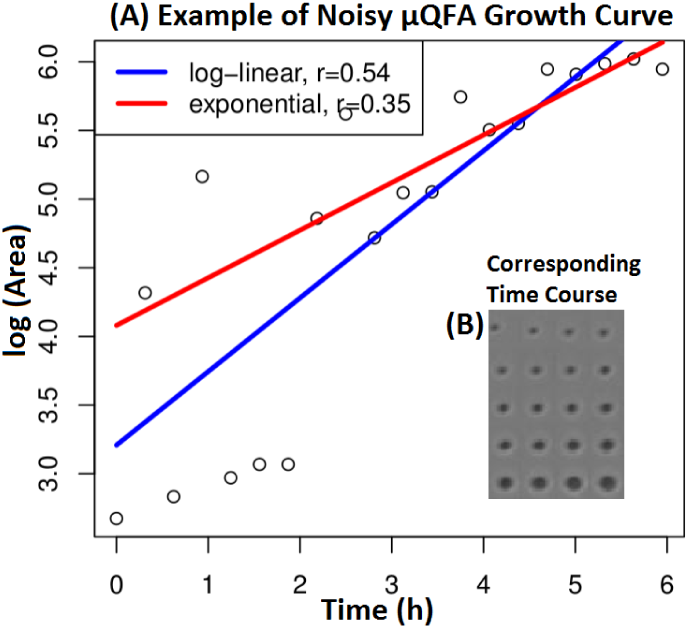
Example *μ*ΟΕΑ growth curve: Modeling the displayed growth curve (A) of *his3*Δ gives the fastest growth rate estimate when using the log-linear model. The corresponding (blurry) time course image (B) as observed under the microscope and fed to the image analysis tool is shown.

Subsequently, even though the log-linear model shows a slightly better fit to the data, (maximum standard error, *σ* =0.09, versus *σ* = 0.25 for the exponential model), the log-linear model places, as highlighted in Figure 14, greater emphasis on early time points which are associated with greater uncertainty. Assuming that area estimates for larger colonies are more precise, the log-linear model is likely to overestimate growth rate. Applying a log-linear regression to the lineage area estimates obtained from the *μ* QFA data results in non-constant residuals across lineage size (Figure 15 (C) and (D)), agreeing with existing literature in that cell growth variability decreases with post-inoculation time (Robinson *et al*., 2001). Figure 15 suggest that the assumption of a log-normal measurement model is unjustified for the analyzed *μ* QFA data.

**Fig. 15:**
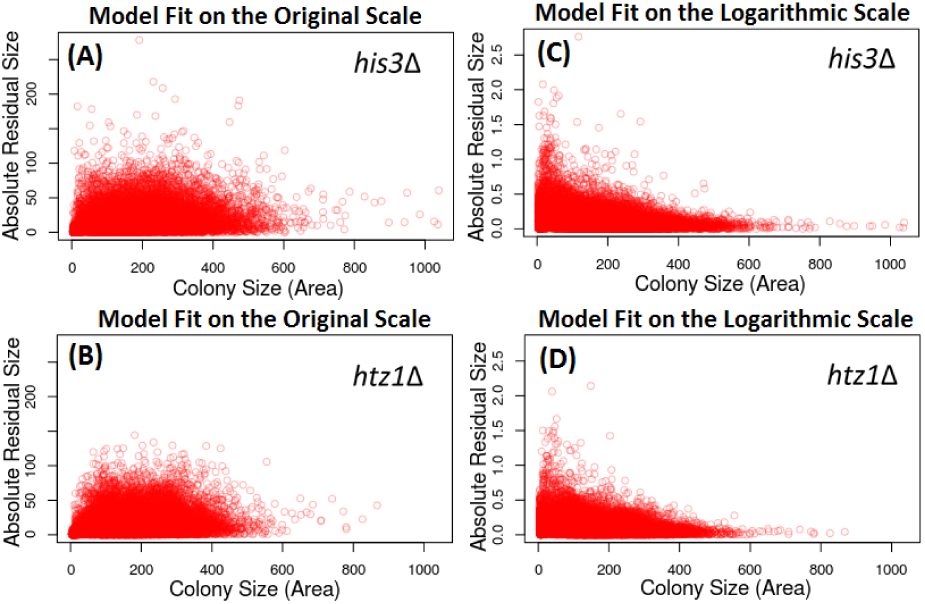
Magnitude of residual values with increasing colony size: Absolute residual size for *his3*Δ (A, C) and *htz1*Δ(B, D) for growth rate estimates using the original exponential model applied using least squares optimization and the log-linear regression model (Figure 13) are plotted against the population size associated with each residual for all *μ* QFA growth curves.

I therefore chose to infer growth rate distributions using the exponential model on the original scale with error also measured on the original scale. Given that error size, even on the original scale, is variable (Figure 15 (A, B)), I chose to confirm inferred parameters with a Bayesian approach by which I can account for an appropriate range of noise within the data using a precision parameter with a wide prior distribution. Bayesian parameter inference carried out using `rjags` (version 4.1.0; Plummer (2016)) confirmed the growth rate densities obtained by frequentist methods (Figure 16).

**Fig. 16:**
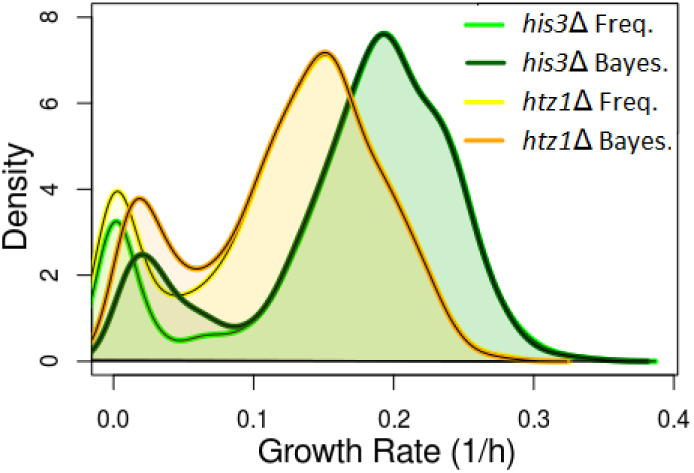
Frequentist versus Bayesian growth rate estimates: Density plots of lineage growth rates of *his3*Δ (green) and *htz1*Δ(yellow) estimated by least squares optimization (Freq.) and Bayesian inference using Gibbs’s sampling (Bayes.) of the exponential growth model are shown.

For lineages with *r* ≥ 0.1, the frequentist point estimates are in good agreement with the expected values of the Bayesian posterior distributions. Frequentist parameter estimation for noisy, non-dividing lineages occasionally resulted in negative growth rates which I set to zero; for Bayesian inference, however, the prior distribution for r sets an initial growth rate constraint of *r* ≥ 0, causing slight discrepancies in growth rate estimates between the two approaches for *r* < 0.1.

Given that the Bayesian approach returns a probability distribution rather than a single point estimate, I used growth rate estimates obtained from the Bayesian inference for the population simulations presented in the next two sections. As expected, the wild-type surrogate *his3*Δ grows faster, whereas *htz1*Δ displays slightly more heterogeneity as indicated by the slightly wider peak in the distribution (Figure 16).

#### 4.2 Current Bayesian discrete stochastic, birth-only inference techniques are unsuited for high-throughput parameter estimation

Because a Bayesian approach was most successful when modeling *μ* QFA deterministically, I tried a Bayesian approach for parameter inference using a discrete/stochastic hybrid model (Figure 13). Several issues, however, keep this implementation from being suitable for inference at a high throughput level:

i. Upon trying a range of tuning parameters for updating sample estimates, I was unable to find a suitable tuning parameter for doing inference on slow-growing growth curves as parameter updates for values close to zero are always attracted to zero. When testing tuning parameters between 001 and 1, the model did not converge to a sensible parameter estimate within 40000000 iterations.
ii. Convergence for growth curves with little to no noise (*σ* < 5) cannot be achieved in a reasonable time frame (< 5 days on a commercially available laptop). With precision approaching infinity, the probability of acceptance approaches zero (Chen *et al*., 2010).
iii. Noise values in the current implementation are constant. No constant value applies to all *μ* QFA growth curves. An implementation where noise is estimated (Wilkinson, 2006) should be considered in the future, but only once convergence for different constant noise values can be achieved on a wide range of growth curves including slow-growing lineages.

Nonetheless, for growth curves with a reasonable amount of noise and growth rate *r* > 0.2, parameter values converge to give a growth rate estimate. For example, the growth rate parameter obtained from the Bayesian discrete stochastic inference (which assumes a normal error measurement model) shown in Figure 17 falls in between the two deterministic parameter estimates (Figure 14) suggesting that when considering inter-lineage stochasticities, a slightly faster growth rate than the one I use in my simulations should be adopted for this lineage.

**Fig. 17:**
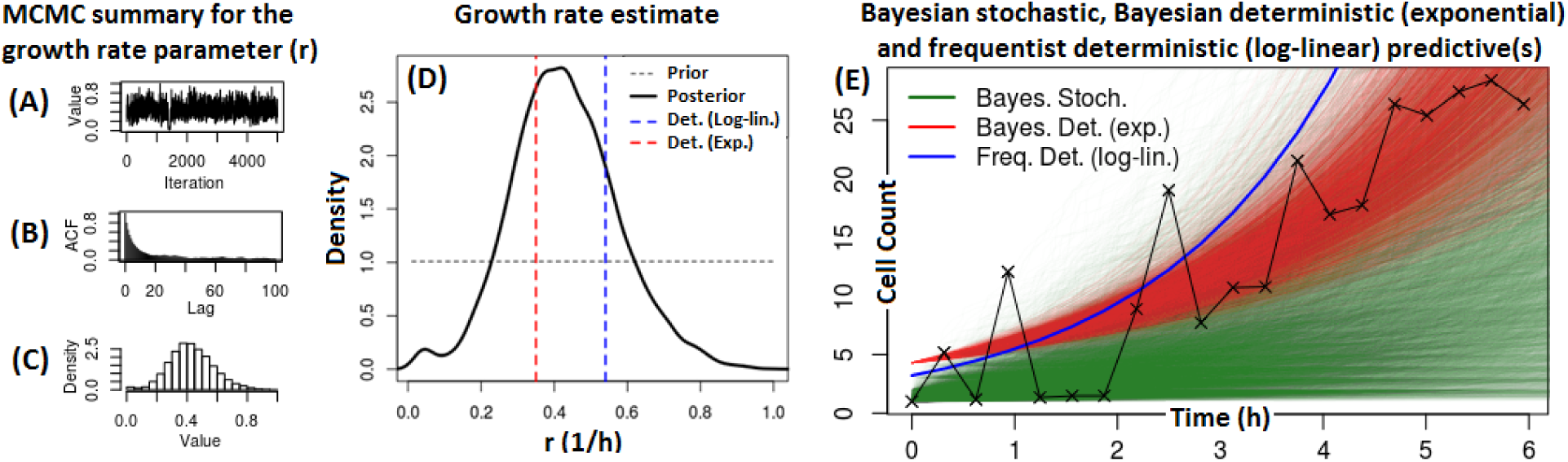
Bayesian stochastic parameter inference accounts for inter-lineage heterogeneities: Bayesian parameter inference outputs when using a hybrid model (Figure 13) on a noisy growth curve (Figure 14) are displayed. Markov Chain Monte Carlo (MCMC) estimates show convergence (A). Auto-correlation tests show limited similarity between subsequent samples (B). Normally distributed parameter estimates are obtained (C). The obtained probability distribution for growth rate *r*, is compared to the deterministic (Det.) estimates (D). 10000 Bayesian stochastic posterior predictives (dark green) are compared to 10000 Bayesian deterministic posterior predictives and the log-linear regression estimate (blue) (E).

#### 4.3 Untrimmed *μ* QFA data confirm the existence of an apparent lag phase at the population level

The *μ* QFA data, which includes non-dividing lineages, confirms the results of Chapter 1, in that all population simulations show a piece-wise linear fit on the log scale. However, population simulations for the *μ* QFA data predict an increase in apparent lag phase duration with increased inoculation size (Figure 18). Growth rate distribution for both *his3*Δ and *htz1*Δ are continuous and have shorter right-hand tails than the distributions assessed in Chapter 1. Contrary to Chapter 1, variability in observed growth rate decreases with increased inoculation size (Figure 19). This confirms the earlier conclusion that an increase in variability with population size is the result of outlying, fast-growing lineages.

**Fig. 18:**
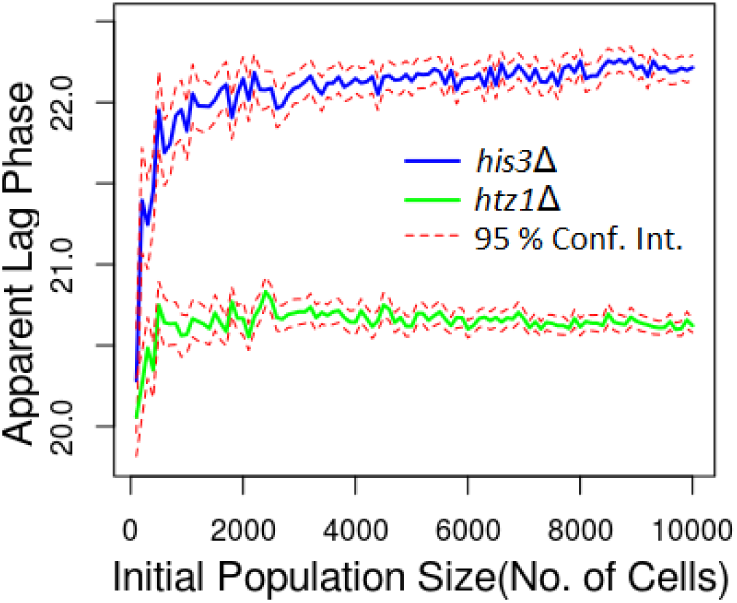
Break point estimates over time in simulations of *his3*Δ and *htz1*Δ: Apparent lag phase duration obtained by fitting Equation 7 to the population simulations generated using Equation 8. Mean estimates along with their 95% confidence intervals (Conf. Int.) of 100 iterations are shown for integer population sizes *n* = [100, 10000] in steps of 100.

**Fig. 19:**
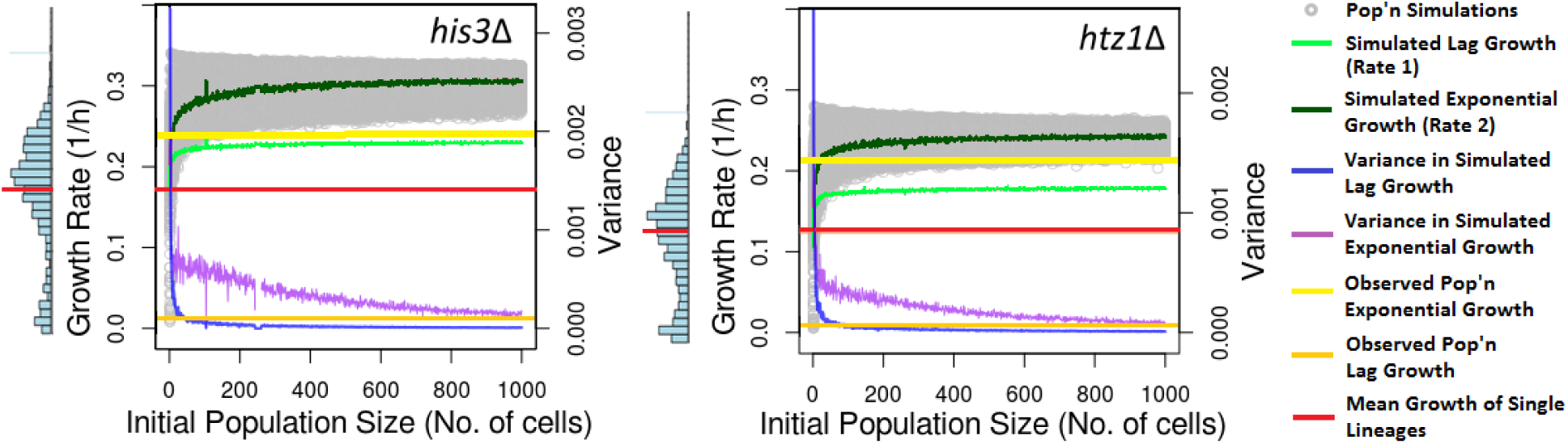
Simulated population growth overestimates observed population growth: Simulated population growth depicted in gray, consisting of 1 - 1000 lineages, is displayed. The mean and variance of 1000 iterations of the two segments of a piece-wise linear regression were calculated and averages for each population start size are displayed in light green and blue for segment 1 (lag) and dark green and purple for segment 2 (exponential). The mean of the two distributions shown vertically on the left of each plot are marked in red. Growth rate distribution scales match those of the main plots. Mean observed population growth for the lag and exponential phase as estimated from Figure 20 are shown in orange and yellow and respectively. The latter estimates are based on mean pin inoculation sizes ranging from 150 to 650 cells.

#### 4.4 *μ* QFA population observations show a lag phase even though single lineage lineage observations do not

As predicted by my simulations, *μ* QFA population observations of each of the pins show a lag phase (Figure 20) even though the corresponding single lineage data provides no evidence for any lag phase. This validates the conclusion from Chapter 1 that an apparent lag phase can arise at the population level as a result of selection. Notably, apparent onset of stationary growth for final observations in Figure 20 must be treated with caution as area estimates are based on two-dimensional observations and cells are likely to be piling on top of each other during final observations resulting in apparent slow growth. Population observations show shorter lag phase duration than predicted by my simulations (Figure 19); this is likely due to small sample sizes in pins which may not capture the full range of single-lineage heterogeneity observed. Pins thus capture a different selection process among single lineages.

**Fig. 20:**
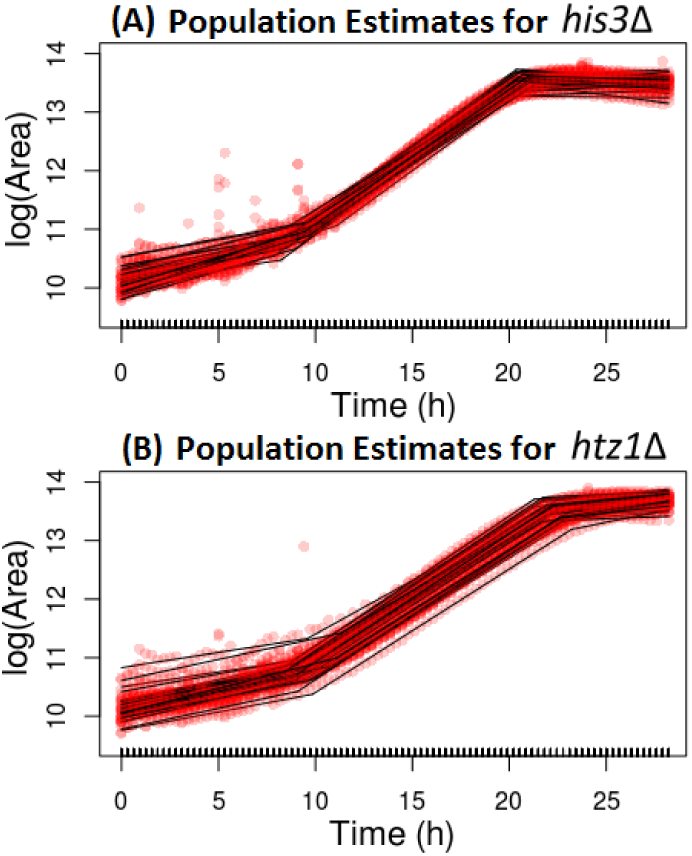
Pin observations of *μ* QFA confirm a lag phase at the population level: Observed population growth for each of the spots (pins) analyzed under the microscope is shown. Total area of cells, counted in the form of dark pixels on a lighter agar background as implemented in `muqfatc`, over time provide population growth curves for each pin observed under the microscope. Estimated cell densities over time are shown in red. Black lines indicate a piece-wise line of best fit for each pin.

#### 4.5 Population simulations overestimate lag growth rate

Population simulations predict slightly faster growth rates for the exponential phase and predict significantly faster growth rates than observed during the lag phase of the population data (Figure 19). Nonetheless, both the simulated and observed population estimates show a significant distinctions between lag and exponential growth whereby exponential population growth surpasses mean growth rates observed at the single lineage level. This again confirms selection of fast-growing lineages driving observed population growth rates.

When sampling from lineage distributions associated with a single pin and comparing the simulations to the population observations, the two estimates match more closely (Figure 21). Although less visible on this scale, simulated lag and exponential growth are significantly different. The predicted distinction in growth rate between the two phases is clearly confirmed at the population level. A predicted lag phase duration of 12.5 h for pin *R*05*C*05 is also confirmed by the true population observations. However, population simulations still significantly overestimate lag growth rates. Figure 19 suggests that mean pin lag growth (orange) does not match the mean growth rate (red) observed among single lineages but captures a much greater number of non-growing or slow-growing lineages possibly missed or discarded by the image analysis. Subsequent exponential growth rates only differ by 0.02 h^−1^.

**Fig. 21:**
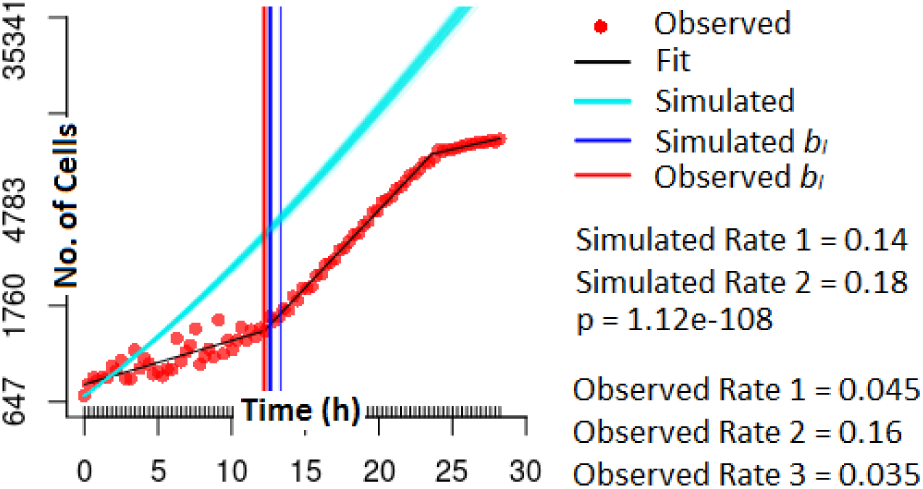
Observed and simulated population growth in pin *R*05*C*05: Observed population growth up to 28 h obtained from *htz1*Δ cells inoculated on solid agar in row 5 column 5 (*R*05*C*05). The black line shows a best-fit piece-wise linear regression with two break points. 1000 population simulations derived from the growth rate estimates from pin *R*05*C*05 and simulated according to Equation 8 with the starting population set to the number of single cells in the pin at time point zero, are overlayed in cyan. Break point estimates for the population simulations (blue) and the first break point in the observed population data (red) are shown. Mean growth rates for all phases are displayed on the right.

### 5 DISCUSSION

*μ* QFA data, unlike the lineage data considered in Chapter1, are best modeled using a normal error measurement model rather than a log-normal one due to high uncertainty in early time points. Choosing a model that accurately captures the validity of the data is absolutely necessary when doing simulations where single outliers significantly influence simulated population growth. A Bayesian approach to parameter inference is preferred since it returns a probability distribution rather than a point estimate and provides greater flexibility in model error measurement. Similar to Chapter1, the discrete stochastic parameter inferences suggests that considering inter-lineage stochasticities provides slightly different parameter estimates. This could be particularly crucial for estimating growth rates of fast-growing lineages and should be taken into consideration once an implementation capable of handling high-throughput single lineage growth data including slow-growing and non-dividing growth curves is achieved. While I here present a birth-only hybrid model for Bayesian parameter inference in R (version 3.3.2; R Core Team, 2016), I propose re-writing the implementation in a compiled programming language to increase computational speed as parameter inference can currently take more than five days to achieve convergence when run on a commercially available laptop. An attempt to enhance the current implementation was made using the Rcpp package (version 0.12.5; Eddelbuettel and Francois, 2011) which allows for C++ code to be integrated into R; however, this only increased computational speed by a negligible amount. Functional programming techniques could be used to run particle filters in parallel (Wilkinson, 2016); an implementation in Scala (Odersky *et al*., 2006), for example, could prove very useful.

Raw image data of *μ* QFA allow for a verification of simulated population growth with true observed population growth for each pin on an agar plate. Observed population growth confirms that a lag phase arises at the population level, even when there is no evidence for a lag phase at the single lineage level. To my knowledge, I present the first attempt at capturing the implications of single lineage growth rate heterogeneity and verifying these at the population level, providing a new level of detail to existing predictive microbiology approaches (Baranyi and Roberts, 1994). The fact that my simulations overestimate lag growth rate at the population level suggests that further work may need to be done to estimate single lineage growth more precisely. However, pin population observations consist of small sample sizes which are unlikely to capture the full range of heterogeneity of strain lineages; thus, to test population observations against simulations, the entire plate as done in QFA (Addinall *et al*., 2011) should be considered in the future. Furthermore, the prediction that the lag phase lengthens with inoculation size has yet to be confirmed for *μ*QFA.

The question of the origin of growth rate heterogeneity addressed at the end of Chapter 1 remains open. Differences in cell age have been shown to result in growth rate heterogeneity at the single lineage level (Ginovart *et al*., 2011). The current literature also suggests that heterogeneity in single lineage growth rate frequently arises as a result of changing environmental conditions and thus continues to persist in populations (Levy *et al*., 2012; Cooper *et al*., 2001; Batchelor *et al*., 1997). Recent work by Van Dijk *et al*. (2015) has shown that even the response to stress is vastly heterogeneous among isogenic lineages. Avraham *et al*. (2013) further show that low-zinc conditions result in continued division of mother cells and complete arrest of daughter cells, demonstrating extreme differences in growth rate at the single cell level. It is thus plausible that growth rate heterogeneity observed at the lineage level translates down to the single cell level. Each reproducing cell may produce offspring according to a given growth rate distribution. While stochastic modeling captures interlineage variability, quantifying selection among individual cells within lineages would require that single cells be isolated and observed at various growth stages, an experimental undertaking which has yet to be done for *S. cerevisiae*.

As demonstrated, growth rate heterogeneity should certainly be measured at a scale smaller than purely population observations in order to fully capture growth rate as a strain phenotype and assess slow-growing and non-dividing subpopulations.

## ACKNOWLEDGEMENT

This work was supported by the supervision of Conor Lawless who proposed the original project and provided continuous guidance along the way, and the supervision of Paolo Zuliani who provided valuable feedback at various project stages. Data for Chapter 1 (Levy *et al*., 2012) was kindly provided by Sasha Levy and Mark Siegal. The discrete stochastic parameter inference outlined in Chapter 1 was done using an (unpublished) implementation kindly provided by Jeremy Revell. Experimental proceedings for Chapter 2 were made possible by David Lydall and his laboratory group who also provided constructive project feedback in response to a presentation I held at one of their internal seminars.

